# Immune Profiling of Dermatologic Adverse Events from Checkpoint Blockade using Tissue Cyclic Immunofluorescence

**DOI:** 10.1101/2023.04.03.535435

**Authors:** Zoltan Maliga, Daniel Y. Kim, Ai-Tram N. Bui, Jia-Ren Lin, Anna K. Dewan, George F. Murphy, Ajit J. Nirmal, Christine G. Lian, Peter K. Sorger, Nicole R. LeBoeuf

**Affiliations:** Laboratory of Systems Pharmacology, Harvard Medical School, Boston, Massachusetts; Harvard-MIT Health Sciences and Technology Program, Harvard Medical School, Boston, Massachusetts; Department of Dermatology, The Center for Cutaneous Oncology, Dana-Farber Cancer Institute and Brigham and Women’s Hospital, Boston, Massachusetts; Department of Dermatology, Vanderbilt University Medical Center, Nashville, Tennessee; Program in Dermatopathology, Department of Pathology, Brigham and Women’s Hospital, Boston, Massachusetts

**Author notes:** **Corresponding Author:** Nicole R. LeBoeuf, MD, MPH, Department of Dermatology, The Centers for Melanoma and Cutaneous Oncology, Dana-Farber Cancer Institute, 375 Longwood Avenue, Boston, MA 02115. These authors contributed equally (Zoltan Maliga, Daniel Y. Kim). **Funding Sources:** U2C-CA233262 and Ludwig Cancer Research. **IRB Approval Status:** This project was deemed exempt by the Mass General Brigham Institutional Review Board. **Conflict of Interest Disclosure:** Mr. Kim is a paid consultant at Verve Therapeutics and SeQure Dx, unrelated to this research. Mr. Kim’s interests were reviewed and are managed by Mass General Brigham in accordance with their conflict-of-interest policies.

**Keywords:** Immune-related adverse events, immunotherapy, immune checkpoint blockade, tissue cyclic immunofluorescence, cytotoxic T-lymphocyte associated protein 4, programmed cell death 1, programmed cell death ligand 1, cutaneous adverse events

## Abstract

In this study, we demonstrate the utility of whole-slide CyCIF (tissue-based cyclic immunofluorescence) imaging for characterizing immune cell infiltrates in immune checkpoint inhibitor (ICI)-induced dermatologic adverse events (dAEs). We analyzed six cases of ICI-induced dAEs, including lichenoid, bullous pemphigoid, psoriasis, and eczematous eruptions, comparing immune profiling results obtained using both standard immunohistochemistry (IHC) and CyCIF. Our findings indicate that CyCIF provides more detailed and precise single-cell characterization of immune cell infiltrates than IHC, which relies on semi-quantitative scoring by pathologists. This pilot study highlights the potential of CyCIF to advance our understanding of the immune environment in dAEs by revealing tissue-level spatial patterns of immune cell infiltrates, allowing for more precise phenotypic distinctions and deeper exploration of disease mechanisms. By demonstrating that CyCIF can be performed on friable tissues, such as bullous pemphigoid, we provide a foundation for future studies to examine the drivers of specific dAEs using larger cohorts of phenotyped toxicity and suggest a broader role for highly multiplexed tissue imaging in phenotyping the immune mediated disease that they resemble.

## Main Text

By inhibiting key immunomodulatory pathways, immune checkpoint inhibitors (ICIs) can have profound anti-tumor effects in cancer patients. However, immune-related adverse events are observed in many patients on ICI therapy, the most frequent of which are dermatologic adverse events (dAEs; 35-50%).^1,2^ dAEs are characterized by diverse reaction patterns, visible clinical specificity, and easily accessed tissues, but deep phenotyping of skin immune toxicity is limited. Histological phenotyping primarily involves immunohistochemistry (IHC), in which serial tissue sections are stained with one of a few antibodies and corresponding signal is scored by a pathologist on a simple numerical scale, commonly 0-4, with 4 indicating strong staining for a specific marker. This approach can be effective in diagnosis or simple prognostication but does not provide the in-depth single cell data necessary to precisely subtype immune cells and characterize mechanisms of activation and engagement with epithelial cells.^3^ Compared to single channel IHC, highly multiplexed tissue imaging has a key advantage: measuring the levels of many antigens at a single cell level makes it possible to precisely determine cell lineage and state.^4,5^ Whole-slide CyCIF (tissue based cyclic immunofluorescence) imaging has the added advantage that it is compatible with clinical histopathology workflows, leverages commercially available antibodies, and is relatively low cost. In CyCIF, iterative rounds of tissue staining, 4-6 channel imaging, and fluorophore inactivation are used to assemble highly multiplexed image at subcellular resolution.^6^ Most applications of CyCIF and other high-plex tissue imaging to date have been performed on tumors with close-packed cells, and it is not clear whether friable tissues such as skin can be as readily analyzed. Here, we demonstrate that CyCIF can be used to profile non-tumor skin across a diverse array of inflammatory cutaneous toxicities caused by ICIs; this highlights its advantages over skin profiling using IHC.

We identified six cases of ICI-induced dAEs from the Skin Toxicities Program at the Dana-Farber and Brigham and Women’s Cancer Center pathology archive, four of which were induced by PD-1 inhibitor monotherapy, one by PD-L1 inhibitor monotherapy, and one by PD-1/CTLA-4 inhibitor combination therapy. These six cases included a range of inflammatory dermatoses that are observed in ICI-treated patients, including lichenoid, bullous pemphigoid, psoriasis, and eczematous eruptions (Supplemental Table 1). The immune and cellular profiles of these six dAE specimens were characterized using both standard IHC (12 antibodies) and six cycle CyCIF (15 antibodies plus a nuclear stain) (Supplementary Table 2 & 3). As an example, Figure 1A and 1B shows staining patterns for an anti-PD-1-induced psoriasis case using 12 IHC antibodies and 15 CyCIF antibodies. The sample was phenotyped using a gating-based approach that distinguishes cells that express or do not express a particular marker (Figure 1C). Unsupervised clustering was then performed to classify distinct cell phenotypes, and cell positions were confirmed through visual inspection of the initial tissue image (Figure 1D & 1E).

**Figure 1.**
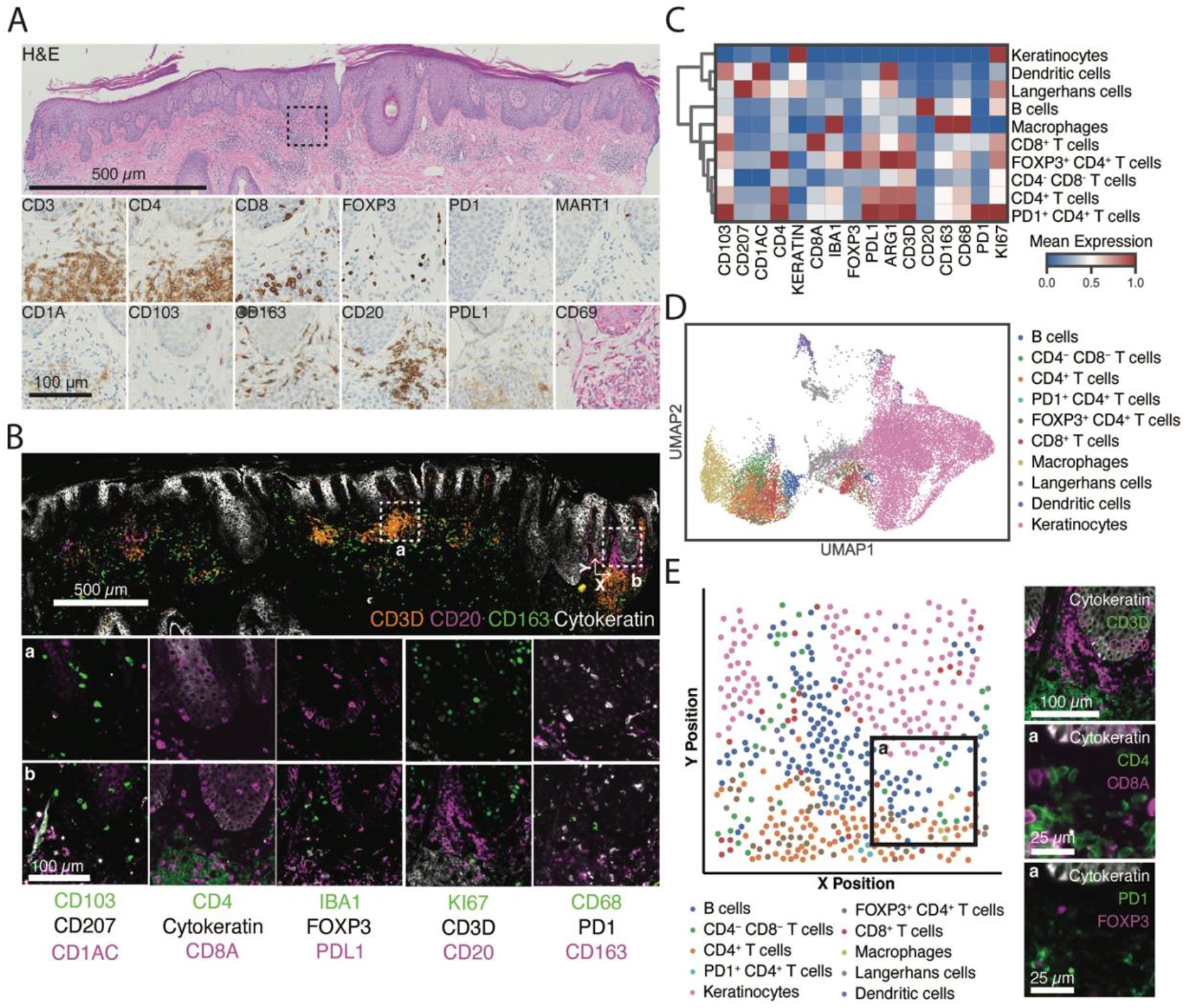
Characterization of PD1-inhibitor induced psoriasis using cyclic immunofluorescence. (A) Full tissue bright-field hematoxylin and eosin-stained image and comparable inset regions (boxed) of immunohistochemistry-stained serial sections. Scale bars, 500 and 100 µm, respectively. (B) Full tissue composite cyclic immunofluorescence (CyCIF) image stained for epidermis (pan-cytokeratin: grayscale), macrophages (CD163: green) and T (CD3D: orange) and B (CD20: magenta) lymphocytes. Not all channels are shown for clarity. Scale bar, 500 µm. Inset, composite images of all CyCIF markers for two lymphoid structures defined by dermal (a) T-cell or (b) mixed B- and T-cell infiltrates. Scale bar, 100 µm. (C) Heatmap showing the mean staining intensity of all markers in each cellular phenotype in tissue sample. (D) Uniform manifold approximation and projection (UMAP) of single-cell data derived from CyCIF labeled by cell type. (E) Scatterplot of cell centroids within inset region (b) colored by cell phenotypes.

After immune phenotyping by CyCIF, we quantified inflammatory cells across all six dAEs and observed total cell counts ranging from 1,615 in the PD-1/CTLA-4 eczematous dermatitis case to 8,242 in the PD-1 psoriasis case (Supplementary Table 4). We then compared the abundance of cells positive for specific markers or combinations of markers with conventional pathology scores (values of 0-4) obtained by pathologist inspection of IHC from 12 serial sections (Figure 2). Across all cases, conventional pathology scores and cell abundances quantified by CyCIF were generally consistent across all four markers that were examined (CD4, CD8, FOXP3, PD-1) (Figure 2A & Figure 2B). However, there were some notable differences between the two methods. For example, immunoreactivity scores for CD4 were 3-4 in all cases, with a score of 3 corresponding to 25%-75% of lymphocytes expressing CD4 and a score of 4 indicating >75% of lymphocytes expressing CD4 (Figure 2A & 2B). In contrast, CyCIF showed that the fraction of CD3D^+^ cells that were also CD4^+^ was most abundant in the eczematous and bullous pemphigoid cases (71-81% of T-cells) and was comparably lower in the psoriasis and lichenoid cases (32-57% of T-cells), which was confirmed through visual inspection of the initial tissue images (Figure 2A-2H). Regarding CD8 expression, the fraction of CD3D^+^ T-cells that were CD8^+^ was most abundant in the two lichenoid cases (58% and 72% of T-cells) and more than 2-fold lower in the PD-1/CTLA-4 eczematous case (22% of T-cells), despite immunoreactivity scores of 3-4 for all samples (Figure 2A & 2B). Upon visual inspection, CD8^+^ expression was confirmed to be highest in the lichenoid cases, which is consistent with prior literature indicating elevated CD8^+^ T-cell infiltrates in lichenoid tissue reactions (Figure 2B & 2C).^7^ With respect to PD-1 levels, standard IHC immunoreactivity scores suggested the greatest PD-1 lymphocyte immunoreactivity in the PD-L1 lichenoid case (immunoreactivity score of 3), but little if any PD-1 expression in all other cases. When profiled using CyCIF, PD-1^+^ T-cell expression was observed in all samples, with greatest abundance in the two lichenoid cases (14 and 21% of T-cells) (Figure 2A & 2B).

**Figure 2.**
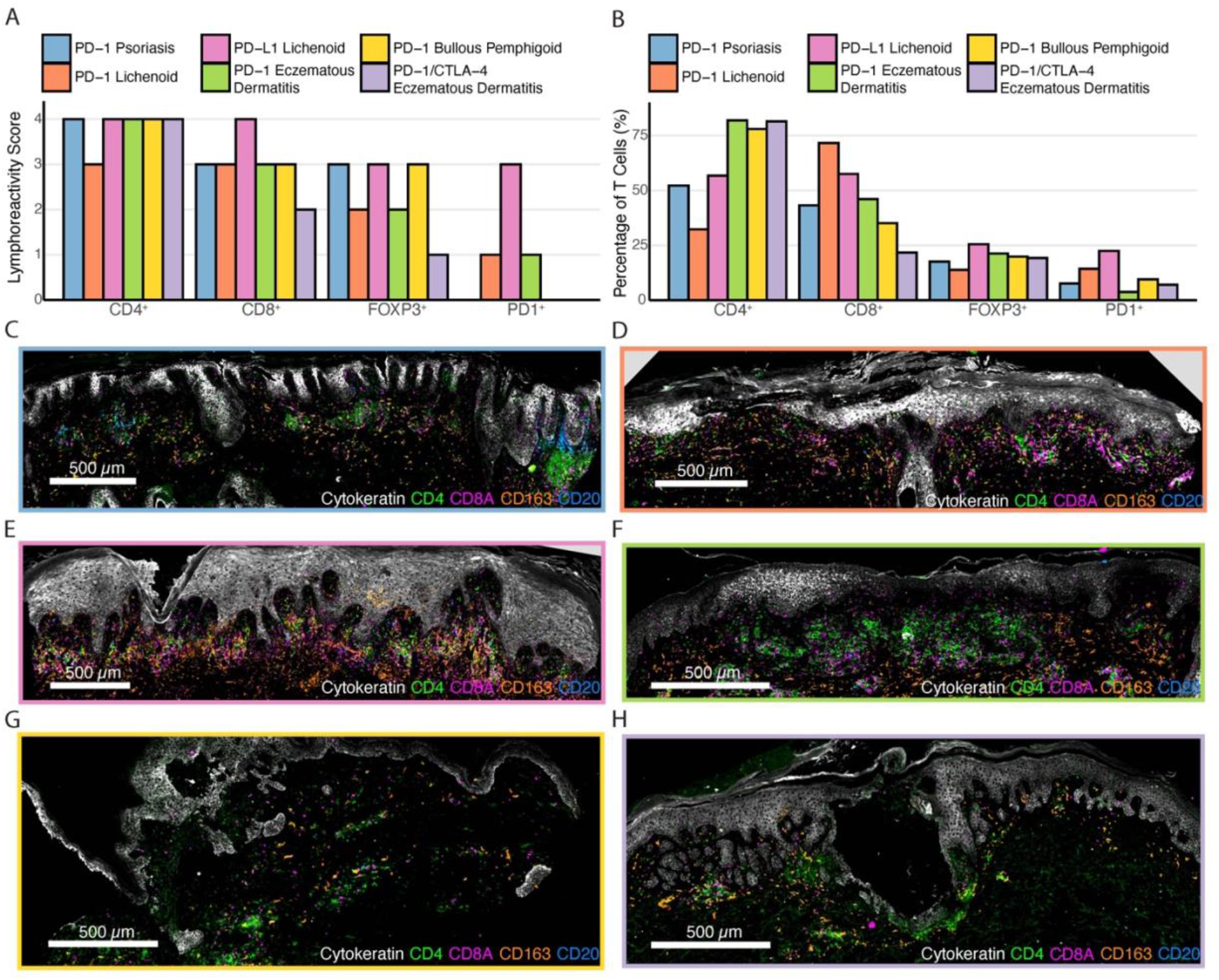
Comparison of conventional histochemical preparations and cyclic immunofluorescence analysis of dermatologic adverse events. (A) Lymphocyte immunoreactivity scores of CD4, CD8, FOXP3, and PD-1 across six dermatologic adverse events using standard immunohistochemistry. (B) Fraction of T-lymphocytes positive for CD4^+^, CD8^+^, FOXP3^+^, and PD1^+^ derived from CyCIF data. Large-field image and composite images for (C) PD-1 inhibitor psoriasis tissue section, (D) PD-1 inhibitor lichenoid tissue section, (E) PD-L1 inhibitor lichenoid tissue section, (F) PD-1 inhibitor eczematous dermatitis tissue section, (G) PD-1 inhibitor bullous pemphigoid tissue section, and (H) PD-1/CTLA-4 inhibitor eczematous dermatitis tissue section.

These data demonstrate that CyCIF can be performed on diverse types of dAEs to provide immune profiles that have greater detail than is feasible with IHC. Moreover, despite general concordance, there were notable differences between human scoring of IHC (on a 0-4 scale) and direct quantitation of immune cell composition in CyCIF using image processing algorithms. These differences are readily explained by the single cell resolution of multiplexed tissue immunofluorescence and by IHC’s reliance on serial sections (each stained by one or two antibodies) which inevitably differ in cell composition. We therefore conclude that CyCIF and similar high-plex imaging methods have the ability to substantially improve our understanding of dAEs. For example, while prior studies have indicated elevated levels of CD4^+^ T-cells in eczematous eruptions and CD8^+^ T-cells in lichenoid morphologies,^7,8^ CyCIF also allows for tissue-level description of spatial patterns, which is a more sensitive way to distinguish phenotypes and draw mechanistic insights than traditional scoring methods Timely recognition and targeted mitigation are clinical priorities as toxicities from ICI can lead to treatment interruption and early mortality. Through a greater understanding of the immune environment in patients with ICI-induced cutaneous toxicities, therapeutic intervention can be optimized and further developed. Using distinct clinical phenotypes of cutaneous toxicity from ICI therapy, we determined that CyCIF can be used to provide single cell, high dimensional imaging of immune infiltrates in skin, including in disorders with friable tissue such as bullous pemphigoid. This is foundational for further study of the drivers of specific dAEs using larger cohorts of phenotyped toxicity. Moroever, success of this pilot study on dAEs suggests a broader role for highly multiplexed tissue imaging in phenotyping the immune mediated diseases they resemble.

