## Supplementary Appendix for "Immune Profiling of Dermatologic Adverse Events from Checkpoint Blockade using Tissue Cyclic Immunofluorescence"

Table of Contents

1. Supplemental Methods...............................................................................Pg. 2-3
2. Supplemental Tables 1-4 ...........................................................................Pg. 4-7
3. References ...................................................................................................Pg. 8

### Methods

#### *Immunohistochemistry Analysis*

We identified six cases of immune checkpoint inhibitor induced dermatologic adverse events from the Skin Toxicities Program at the Dana-Farber/Brigham and Women's Cancer Center pathology archive. Standard immunohistochemistry staining for CD4, CD8, FOXP3, and PD-1 were performed by the Brigham and Women's Hospital Longwood Core Laboratory in Boston, MA. Lymphocyte immunoreactivity was then scored by two independent pathologists on a scale of 0-4, with 4 indicating a high number of lymphocytic immunoreactive cells. In the case of disagreement, a third pathologist reported on the score.

#### *Cyclic Immunofluorescence (CyCIF) Sample Preparation and Analysis*

CyCIF imaging of tissues was performed manually at room temperature, unless otherwise noted, by iterative rounds of tissue staining, imaging and photochemical fluorochrome inactivation, as previously described.[1] Tissues were stained overnight at 4˚C, or for 1 hour at room temperature for secondary antibody conjugates, in a dark, humidified chamber with antibodies diluted in Odyssey Blocking Buffer (Licor) supplemented with 2 µg/ml Hoechst 33520 (Bio-Rad, &), rinsed for 30 minutes in room-temperature phosphate buffered saline (PBS) and mounted with 20% glycerol in PBS using a 22x60mm coverslip. Stained tissues were imaged using a Cytefinder (Rarecyte, Inc. Seattle, WA) automated microscope using a 40x (NA=0.6) objective. After imaging, mounted slides were soaked in room-temperature PBS to detach coverslips, immersed in PBS supplemented with 3% H2O2 and 20 mM NaOH and illuminated. After 1 hour, slides were rinsed twice in PBS in preparation for the next staining cycle. Six rounds of iterative staining and imaging were performed on each sample. Image processing, cell segmentation and thresholding was performed using MCMICRO and scimap.[2,3]

#### *Gating and Phenotyping Strategy*

Cell phenotypes were defined by manual gating each tissue sample in the following sequence: keratinocytes (Cytokeratin$^{+}$), B cells (CD20$^{+}$), Langerhans cells (CD207$^{+}$), dendritic cells (CD1AC$^{+}$), macrophages (CD163$^{+}$), CD4$^{-}$ CD8$^{-}$ T cells (CD4$^{-}$ CD8$^{-}$ CD3D$^{+}$), CD8$^{+}$ T cells (CD8$^{+}$ CD3D$^{+}$), CD4$^{+}$ T cells (CD4$^{+}$ CD3D$^{+}$), FOXP3$^{+}$ CD4$^{+}$ T cells (FOXP3$^{+}$ CD4$^{+}$ CD3D$^{+}$), and PD1$^{+}$ CD4$^{+}$ T cells (PD1$^{+}$ CD4$^{+}$ CD3D$^{+}$).

**Supplemental Table 1. Clinical and histologic features of six cases of immune checkpoint inhibitor (ICI) induced dermatologic adverse events (dAEs).**

| Age | Sex | Malignancy | Drug Target | Prior Therapies | Clinical Morphology | Histologic Morphology |
|---|---|---|---|---|---|---|
| 62 | Male | Lung | PD-1 (Pembrolizumab) | Carboplatin + Pemetrexed | Psoriasis | Psoriasis |
| 51 | Female | Cervical | PD1 (Pembrolizumab) | Carboplatin + Paclitaxel + Bevacizumab<br><br>Cisplatin + Radiotherapy<br><br>Cisplatin + Bevacizumab | Lichenoid | Erythema multiforme |
| 83 | Male | Lung adenocarcinoma | PD1 (Nivolumab) | Carboplatin + Pemetrexed | Bullous Pemphigoid | Bullous Pemphigoid |
| 67 | Male | Lung adenosquamous carcinoma | PD-L1 (Atezolizumab) | Cisplatin + Gemcitabine | Psoriasis | Lichenoid |
| 68 | Female | Lung adenocarcinoma | PD1 (Pembrolizumab) | Cisplatin + Pemetrexed<br><br>Ipilimumab + Pembrolizumab | Papular eczema | Spongiotic dermatitis |
| 53 | Male | Melanoma | CTLA-4/PD-1 (Ipilimumab & Nivolumab) | Dabrafenib + Trametinib | Grover’s/Papular eczema | Spongiotic dermatitis |

**Supplemental Table 2. IHC antibody panel for immunophenotyping of dAEs.**

| Target | Target Protein | Catalog Number | Vendor | Clone | Species | Isotype | RRID | Function |
|---|---|---|---|---|---|---|---|---|
| CD1 | CD1 | LS-C122829 | LSBio | RIV12 | mouse | IgG1 | AB_10802995 | Dendritic cells |
| CD103 | CD103 | ab129202 | Abcam | EPR4166(2) | rabbit | IgG | AB_11142856 | Tissue resident memory T lymphocytes |
| CD163 | CD163 | MA5-11458 | Thermo Fisher | 10D6 | mouse | IgG1 | AB_10982556 | Macrophages, M2 |
| CD20 | CD20 | ab78237 | | EP459Y | rabbit | IgG | AB_1640323 | B lymphocytes |
| CD3 | CD3E | ab5690 | Abcam | polyclonal | rabbit | IgG | AB_305055 | T lymphocytes |
| CD4 | CD4 | NBP1-19371 | Novus | polyclonal | rabbit | IgG | AB_1641682 | Helper T lymphocytes |
| CD69 | CD69 | NBP1-51607 | Novus | 8B6 | mouse | IgG1 | AB_11009776 | Tissue resident memory T lymphocytes |
| CD8A | CD8A | ab101500 | Abcam | SP16 | rabbit | IgG | AB_10710024 | Cytotoxic T lymphocytes |
| FOXP3 | FOXP3 | ab10901 | Abcam | polyclonal | rabbit | IgG | AB_297559 | Regulatory T lymphocytes |
| MART 1 | MART 1 | 917901 | Biolegend | M2-7C10 | mouse | IgG2b | AB_2565202 | Melanocytes |
| PD-1 | PD-1 | ab52587 | Abcam | NAT105 | mouse | IgG1 | AB_881954 | Activated or exhausted T lymphocytes |
| PD-L1 | PD-L1 | 13684S | Cell Signaling | E1L3N | rabbit | IgG | AB_2687655 | Immune checkpoint |

### Supplemental Table 3. CyCIF antibody panel for immunophenotyping of dAEs.

| Cycle Number | Antibody Name | Target Protein | Vendor | Catalog Number | Clone | Fluorophoe | RRID | Function |
|---|---|---|---|---|---|---|---|---|
| 1 | CD103 | CD103 | Cell Marque (Millipore Sigma) | 437R-15 | EP206 | anti-rabbit IgG Alexa Fluor 488 | AB_2884943 | Tissue resident memory T lymphocytes |
| 1 | Langerin | Langerin | R&D Systems | AF2088-SP | polyclonal | anti-goat IgG Alexa Fluor 555 | AB_355143 | Langerhans Cells |
| 1 | CD1c | CD1c | Abcam | ab156708 | OTI2F4 | anti-mouse IgG Alexa Fluor 647 | AB_2889187 | Dendritic Cells |
| 1 | CD1a | CD1a | Abcam | ab201337 | C1A/711 | anti-mouse IgG Alexa Fluor 647 | AB_2889186 | Dendritic Cells |
| 1 | Rabbit IgG | Rabbit IgG | Thermo Fisher Scientific (eBioscience) | | polyclonal | | AB_143165 | |
| 1 | Goat IgG | Goat IgG | Thermo Fisher Scientific (eBioscience) | | polyclonal | | AB_2535853 | |
| 1 | Mouse IgG | Mouse IgG | Thermo Fisher Scientific (eBioscience) | | polyclonal | | AB_10564125 | |
| 2 | CD4 | CD4 | R&D Systems | FAB8165G | polyclonal | Alexa Fluor 488 | AB_2728839 | Helper T lymphocytes |
| 2 | Cytokeratin (pan) | Cytokeratin (pan) | Thermo Fisher Scientific (eBioscience) | 41-9003-80, 41-9003-82 | AE1/AE3 | eFluor 570 | AB_11218704 | Epidermis, ducts |
| 2 | CD8a | CD8a | eBioscience | 50-0008-82 | AMC908 | eFluor 660 | AB_2574149 | Cytotoxic T lymphocytes |
| 3 | IBA1 | IBA1 | Abcam | ab195031 | EPR6136(2) | Alexa Fluor 488 | AB_2889157 | Myeloid activation |
| 3 | FOXP3 | FOXP3 | eBioscience | 41-4777-80, 41-4777-82 | 236A/E7 | eFluor 570 | AB_2573609 | Regulatory T lymphocytes |
| 3 | PD-L1 | PD-L1 | Cell Signaling Technology | 15005 | E1L3N | Alexa Fluor 647 | AB_2728832 | Immune checkpoint |
| 4 | ARG1 | ARG1 | Cell Signaling Technology | 66297 | D4E3M | Alexa Fluor 488 | AB_2799705 | Immune checkpoint |
| 4 | CD3 | CD3D | Abcam | ab208514 | EP4426 | Alexa Fluor 555 | AB_2728789 | T lymphocytes |
| 4 | CD20 | CD20 | eBioscience | 50-0202-80, 50-0202-82 | L26 | eFluor 660 | AB_11151691 | B lymphocytes |
| 5 | CD163 | CD163 | Abcam | ab218293 | EPR14643-36 | Alexa Fluor 488 | AB_2889155 | Macrophages, M2 |
| 5 | CD68 | CD68 | Cell Signaling Technology | 79594, 79594S | D4B9C | Phycoerythrin | AB_2799935 | Macrophages, M1 |
| 5 | PD-1 | PD-1 | Abcam | ab201825 | EPR4877(2) | Alexa Fluor 647 | AB_2728811 | Activated or exhausted T lymphocytes |
| 6 | KI67 | KI67 | eBioscience | 41-5699-80 | 20Raj1 | eFluor 570 | AB_11220088 | Proliferating cells |

**Supplemental Table 4. Inflammatory cell phenotype counts for six ICI-induced dAEs using CyCIF.**

| | PD-1 Psoriasis | PD-1 Lichenoid | PD-1 Eczematous Dermatitis | PD-1 Bullous Pemphigoid | PD-L1 Psoriasis | PD-1/CTLA-4 Eczematous Dermatitis |
|---|---|---|---|---|---|---|
| **Cells, No. (%)** | 8,242 | 6,007 | 4,737 | 2,242 | 5,512 | 1,615 |
| Dendritic cells | 627 (7.6%) | 164 (2.7%) | 534 (11.3%) | 381 (17.0%) | 71 (1.3%) | 274 (17.0%) |
| Langerhans cells | 1332 (16.2%) | 262 (4.4%) | 89 (1.9%) | 157 (7.0%) | 157 (2.8%) | 149 (9.2%) |
| Macrophages | 1538 (18.7%) | 1442 (24.0%) | 989 (20.9%) | 269 (12.0%) | 1934 (35.1%) | 300 (18.6%) |
| B cells | 391 (4.7%) | 227 (3.8%) | 89 (1.9%) | 43 (1.9%) | 198 (3.6%) | 196 (12.1%) |
| T cells | 4354 (52.8%) | 3912 (65.1%) | 3033 (64.1%) | 1392 (62.1%) | 3152 (57.2%) | 696 (43.1%) |
| $CD4^+$ T cells | 1366 (31.4%) | 340 (8.7%) | 1282 (42.3%) | 623 (44.8%) | 693 (22.0%) | 389 (55.9%) |
| $FOXP3^+$ $CD4^+$ T cells | 543 (12.5%) | 285 (7.3%) | 534 (17.6%) | 214 (15.4%) | 631 (20.0%) | 116 (16.7%) |
| $PD1^+$ $CD4^+$ T cells | 65 (1.5%) | 123 (3.1%) | 82 (2.7%) | 78 (5.6%) | 70 (2.2%) | 20 (2.9%) |
| $CD8^+$ T cells | 1342 (30.8%) | 2586 (66.1%) | 925 (30.5%) | 305 (21.9%) | 1455 (46.2%) | 99 (14.2%) |
| $CD4^-$ $CD8^-$ T cells | 1038 (23.8%) | 578 (14.8%) | 210 (6.9%) | 172 (12.4%) | 303 (9.6%) | 72 (10.3%) |
